# A phased chromosome-level genome and full mitochondrial sequence for the dikaryotic myrtle rust pathogen, *Austropuccinia psidii*

**DOI:** 10.1101/2022.04.22.489119

**Authors:** Richard J Edwards, Chongmei Dong, Robert F Park, Peri A Tobias

## Abstract

The fungal plant pathogen *Austropuccinia psidii* is spreading globally and causing myrtle rust disease symptoms on plants in the family Myrtaceae. *A. psidii* is dikaryotic, with two nuclei that do not exchange genetic material during the dominant phase of its life-cycle. Phased and scaffolded genome resources for rust fungi are important for understanding heterozygosity, mechanisms of pathogenicity, pathogen population structure and for determining the likelihood of disease spread. We have assembled a chromosome-level phased genome for the pandemic biotype of *A. psidii* and, for the first time, show that each nucleus contains 18 chromosomes, in line with other distantly related rust fungi. We show synteny between the two haplo-phased genomes and provide a new tool, ChromSyn, that enables efficient comparisons between chromosomes based on conserved genes. Our genome resource includes a fully assembled and circularised mitochondrial sequence for the pandemic biotype.

## Introduction

Originating in South America, the plant fungal pathogen, *Austropuccinia psidii* (G. Winter) Beenken *comb. nov*, has been spreading globally and causing myrtle rust infection on species of the family Myrtaceae (Carnegie and Pegg 2018). Most of the global spread has been caused by the pandemic biotype with detection in Hawaii since 2005, and in Australia and New Zealand since 2010 and 2017 respectively (Carnegie and Pegg 2018). As a new encounter pathogen for an expanding global species list (Soewarto *et al*. 2019), *A. psidii* has caused dramatic declines in the health of trees and shrubs despite no apparent co-evolution (Tobias *et al*. 2015). Notably in Australia, a centre of diversity for Myrtaceae, with around 1,700 of the more than 5,500 global species (Wilson, 2011), once common species are now listed as critically endangered (Berthon et al. 2018, Fensham et al. 2020) and at least sixteen rainforest trees are predicted to be extinct in the wild within a generation due to myrtle rust (Fensham and Radford-Smith 2021). Alarmingly, species such as *Lenwebbia* sp. Main Range (P.R.Sharpe+ 4877), have been listed as critically endangered before being formally taxonomically described (NSW Threatened Species Scientific Committee 2019).

*Austropuccinia psidii* predominately exists in the asexual dikaryotic urediniospore stage of the life cycle, with two separate haploid nuclei (Tobias *et al*. 2021). During the dominant life-cycle state of clonal reproduction on a living host, the two nuclei do not exchange genetic material, hence the two genomic compartments are known to develop mutations, structural variants and high levels of heterozygosity (Schwessinger *et al*. 2020). Understanding infection mechanisms and life-cycle traits therefore relies on determining the genetic components within the two separate haploid nuclei. Specifically, mating strategies, in obligate biotrophs of the Puccinomycotina, cannot be clearly deciphered without full dikaryon genome assemblies, where the + and – nuclei are required for sexual compatibility (Raper 1960; Coelho *et al*. 2017).

A key goal for researchers of all eukaryotic organisms is the fully phased diploid genome with telomere to telomere (T2T) chromosomes. The complete gapless, human genome was recently published after decades of intensive collaborative efforts to achieve these goals (Sergey *et al*. 2022). A comprehensive genome for the pandemic biotype was recently published using PacBio (Pacific Biosciences of California, Inc.) RSII and Sequel sequencing technology (Tobias *et al*. 2021). The genome provided new knowledge on the large composition of transposable elements, greater than 90%, unusually large telomeres, and determined that the haploid size was around 1 Gbp. Scaffolding the genome, however, remained problematic due to error-prone early generation PacBio sequence reads likely confounding the assembly and scaffolding software. Here, we present a phased chromosome-level genome assembly and make available a complete circularised sequence of the mitochondrial genome for the pandemic biotype of *A. psidii*. We show for the first time that *A. psidii* has 18 haploid chromosomes, thereby conforming to published karyotypes for other rust species (Boehm and Bushnell 1992; Boehm *et al*. 1992a) and we make available a new tool for efficient high-level synteny analysis between genomes, ChromSyn (https://github.com/slimsuite/chromsyn).

## Methods

### DNA sequenced on PacBio Sequel II

Our research group previously obtained high molecular weight (HMW) DNA from the pandemic biotype of *A. psidii* (Tobias et al. 2021) and sequenced the long reads using PacBio RSII and Sequel technology. The data, along with reads from Hi-C technology (van Berkum et al. 2010) were used to assemble a one gigabase pair (Gbp) genome. To improve the scaffolding on the previous assembly with more accurate sequencing technology, we sent 16 micrograms of the same sample of HMW DNA to The Australian Genome Research Facility (AGRF) at the University of Queensland, Brisbane, Australia for PacBio Sequel II HiFi sequencing.

### Genome assembly and phasing

We obtained 2,306,603 high quality HiFi reads (Q20) with mean bp length of 14,920 Q20, and median read quality Q33. We assembled the data using 33 X read coverage on the University of Sydney high performance compute cluster, running PBS-Pro (v 13.1.0.160576), with Hifiasm software (v 0.15.4) (Cheng *et al*. 2021) and incorporated 80 GB of raw Hi-C data from our earlier assembly (Tobias *et al*. 2021) to help phase the genomes. Apart from these additional parameters, -t24 -l3, we used default settings and the assembly completed in five hours using 80 GB of memory. The two assembly outputs were around 1 Gbp each, confirming the large genome size determined previously.

### Genome scaffolding with Hi-C data

Phased genome outputs, from the Hifiasm assembly that incorporated Hi-C data, were then independently scaffolded by using the Aiden Lab pipelines (https://github.com/aidenlab). Firstly the files were run through the Juicer pipeline (v 1.6) (Durand *et al*. 2016b) with default parameters. The final output from Juicer was used with the 3D-DNA pipeline (v 180922) (Dudchenko *et al*. 2017) with the following parameters ‘-m haploid --build-gapped-map --sort-output’. After manually curating the assemblies locally within the Juicebox visualisation software (v 1.11.08 for Windows) (Durand *et al*. 2016a) the revised assembly file was resubmitted to the 3D-DNA post review pipeline with the following parameters ‘--build-gapped-map --sort-output’ for final assembly and fasta files.

### mtDNA data extraction and assembly

Initial screening of the two haploid genomes, hereon arbitrarily designated (APSI for *<u>A</u>ustropuccinia <u>psi</u>dii*) **hap1** and **hap2**, identified scaffolds corresponding to partial mitochondrial genomes in addition to a possible full-length mtDNA assembled into unplaced hap1 scaffold 21. HiFi reads were mapped to the full hap1 assembly with minimap v2.22 (Li 2018) and reads that mapped to mtDNA scaffolds, or the mtDNA region of scaffold 21, were extracted using samtools v1.15 (Li *et al*. 2009). mtDNA reads were assembled with Hifiasm v0.16.1 (Cheng *et al*. 2021) and a 94.5 kb circular contig representing the mtDNA was identified. Error-correction was performed by mapping reads back on to a double-copy draft mtDNA using Minimap v2.22 (Li 2018) and polishing with HyPo v1.0.3 **(https://github.com/kensung-lab/hypo)**. A very uneven depth profile was identified, raising concerns that the assembled reads included both pure mtDNA and nuclear mitochondrial insertions (NUMTs). A subset of “pure” mtDNA reads was extracted by searching the double-length draft mtDNA onto all 10+ kb HiFi reads and extracting all reads with 99%+ read coverage by mtDNA using GABLAM v2.30.5 (Davey *et al*. 2007) wrapping BLAST+ v2.11.0 (Altschul *et al*. 1990). This subset was reassembled with Hifiasm and a new 94.5 kb circular contig representing the mtDNA was identified. A double-copy version of this contig was polished with Hypo v1.0.3 using only the “pure” 10 kb reads and re-circularised to have the same starting position as published *A. psidii* mtDNA sequence NC_044121.1 (de Almeida *et al*. 2021).

### Post-scaffolding contamination removal and cleanup

mtDNA-rich scaffolds were identified with NUMTFinder v0.5.1 **(https://github.com/slimsuite/numtfinder)** (Edwards *et al*. 2021) and removed from the assembly. A vecscreen search for adaptors and vecotrs with UniVec, using Diploidocus v1.1.3 (Chen *et al*. 2022), identified an additional scaffold that was pure PacBio synthetic construct, which was removed. Unplaced scaffold 25 contained an 87 bp hit to the PacBio adaptor sequences and was split into two scaffolds, removing the 87 bp fragment. The APSIMTv2 mtDNA sequence was then added to each haploid genome, which were filtered for low-quality sequences and false duplications using Diploidocus v1.1.3 (Chen *et al*. 2022). All junk and quarantine scaffolds were discarded, and any unplaced (non-chromosome-level) scaffolds were de-scaffolded into contigs prior to a second round of Diploidocus cleanup. The final set of core and repeat scaffolds were designated as the version two (v2) assembly (hap1, hap2 and mt). The genome assemblies have been submitted to the National Centre for Biological Information (NCBI) at these accessions; JALGQZ000000000 and JALGRA000000000. The APSI_hap1_v2 genome incorporates the fully circularised mitochondrial DNA sequence as the final scaffold.

### Genome quality assessment

Functional completeness of each haploid genome was estimated using BUSCO v5.2.2 (Simão *et al*. 2015) (implementing BLAST+ v2.11.0 (Altschul *et al*. 1990), HMMer v3.3 (Eddy 2010; Potter *et al*. 2018), MetaEuk v20200908 (Rhie *et al*. 2020), Prodigal v2.6.3 (Levy Karin *et al*. 2020) and SEPP v4.3.10 (Mirarab *et al*. 2011) in MetaEuk mode with the basidiomycota_odb10 lineage database. BUSCO predictions were compiled and checked for consistency using BUSCOMP v1.0.1 (Stuart *et al*. 2021). Raw HiFi read kmer completeness and assembly base-error rates were estimated using Merqury v20200318 (Mirarab *et al*. 2011)(implementing BEDTools v2.27.1(Quinlan and Hall 2010), Meryl v20200313, R v3.6.3 (R Development Core Team 2011)and Samtools v1.10 (Li *et al*. 2009) with *k*=21. Estimated copy numbers for BUSCO Complete and Duplicated genes, predicted effector genes, and 100 kb sliding windows, were calculated with DepthKopy v1.0.3 (Chen *et al*. 2022).

### Chromosome synteny analysis

Chromosome synteny analysis was performed using single-copy “Complete” genes from the BUSCO genome completeness runs. Synteny blocks between pairs of assembly were determined collinear runs of matching BUSCO gene identifications. For each assembly pair, BUSCO genes rated as “Complete” in both genomes were ordered and oriented along each chromosome. Synteny blocks were then established as sets of BUSCO genes that were collinear and uninterrupted (*i*.*e*. sharing the same order and strand) in both genomes, starting at the beginning of the first BUSCO gene and extending to the end of the last BUSCO gene in the block. We note that local rearrangements and breakdowns of synteny between BUSCO genes will not be identified and may be falsely marked as syntenic within a synteny block.

Chromosome synteny was then visualised by arranging the chromosomes for each assembly in rows and plotting the synteny blocks between adjacent assemblies. In each case, the chromosome order and orientation were set for one “focus” assembly, and the remaining assemblies arranged in order to maximise the clarity of the synteny plot, propagating out from the focal genome. For each chromosome, the “best hit” in its adjacent assembly was established as that with the maximum total length of shared synteny blocks, and an anchor point established as the mean position along the best hit chromosome of those synteny blocks. Where the majority of synteny was on the opposite strand, the chromosome was reversed and given an “R” suffix in the plot. Chromosomes were then ordered according to their best hits and, within best hits, the anchor points. Synteny blocks sharing the same orientation were plotted blue, whilst inversions are plotted in red. Code for establishing and visualising chromosome synteny based on shared BUSCO genes has been publicly released as a new tool, ChromSyn, under a GNU General Public License v3.0 and is available on GitHub **(https://github.com/slimsuite/chromsyn)**. Synteny of the two *A. psidii* haploid genomes was analysed with ChromSyn v0.9.3 and further comparisons were also made against the recently published fully phased, chromosome-level wheat leaf rust genome, *Puccinia triticina* (Pt76) (Duan *et al*. 2022). We also compared synteny against the previous *A. psidii* genome assembly (Tobias *et al*. 2021).

### Telomere prediction

Terminal telomeric repeats were predicted using an extension of FindTelomeres **(https://github.com/JanaSperschneider/FindTelomeres)** as implemented by Diploidocus v1.2.0 (Chen *et al*. 2022). Briefly, this searches chromosome termini with degenerate canonical telomere sequences (C{2,4}T{1,2}A{1,3} at the 5’ end, and T{1,3}A{1,2}G{2,4} at the 3’ end), flagging a telomere when at least 50% of the terminal 50 bp match the telomeric repeats. Telomeres were extended in 50 bp sliding windows until the repeat match drops below 50%. Telomere repeat sequences were also predicted using the Telomere Identification toolKit (tidk) v0.1.5 **(https://github.com/tolkit/telomeric-identifier)**. Telomeric repeat windows were first identified with tidk search using the repeat unit AACCCT. Telomere plotting was implemented in ChromSyn, plotting Diploidocus telomeres as central black circles and tidk windows with a repeat number of 50+ as blue circles.

### Effector search with version one genome annotations

Effectors are secreted by pathogens to manipulate susceptible host plants and initiate compatible interactions for infection (Lo Presti *et al*. 2015). They are generally small, secreted proteins and are of primary interest due to their key role in recognition in resistant plants and pathogenicity in susceptible plants (Jaswal *et al*. 2020). We mapped the predicted 671 effector coding sequences, annotated for version one (v1) of the pandemic genome (Tobias *et al*. 2021), to each of the current haploid genomes using BBMap (v37.98) (Bushnell 2014). Specifically, the full set of effector sequences were mapped to hap1 and to hap2, independently, with the mapPacBio script and all defaults. Bedtools (v 2.29.2) (Quinlan and Hall 2010) was used to obtain the genome co-ordinates on scaffolds to visualise locations on chromosomes and to determine proximity to telomeric regions.

## Results

### *Austropuccinia psidii* has 18 haploid chromosomes

Phased assembly of combined (33X) HiFi and Hi-C reads with Hifiasm, followed by additional Hi-C scaffolding, yielded two haploid assemblies of approximately 1 Gbp each, named hap1 and hap2 (Table 1). Each contained 18 chromosome-sized scaffolds visible within the Hi-C contact map (Figure 1). Scaffolds are numbered and ordered by size, hence scaffold 1 is the largest at ∼85 Mb and scaffold 18 is the smallest at ∼25 Mb. Thirteen hap1 and twelve hap2 scaffolds are telomere to telomere, indicating likely full-length chromosomes (Table 1 and Figure 2). Comparison to the wheat leaf rust (strain Pt76) genome (Duan *et al*. 2022) shows the same haploid chromosome count of *n*=18, with many of the chromosomes presenting a dominant syntenic relationship with a single myrtle rust scaffold, despite the significant enlargement in chromosome size (Figure 2). Overall, chromosome-sized scaffolds account for 98.5% of hap1 and 99.3% of hap2. BUSCO v5 analysis (basidiomycota_odb10 lineage, MetaEuk mode) predicted gene completeness levels of 90.7% (hap1) and 91.5% (hap2), with only twenty complete BUSCO genes from hap1 and one from hap2 not found on the chromosome scaffolds (Figure 2? 3?).

**Table 1.** Basic genome statistics comparing the v1 *Austropuccina psidii* genome (Tobias *et al*. 2021) here called APSI_hap1_v1 against the current genome assemblies APSI_hap1_v2 (grey).

| Genome | SeqNum<sup>1</sup> | TotLength | MinLength | MaxLength | N50 | L50 | GC |
| --- | --- | --- | --- | --- | --- | --- | --- |
| APSI_hap1_v1 | 66 (3,817) | 1,018,398,822 | 1,003 | 124,273,364 | 56,243,252 | 7 | 33.8 |
| APSI_hap1_v2 | 104 (332) | 1,016,163,381 | 16,158 | 83,706,672 | 67,280,866 | 7 | 33.85 |
| Chromosomes | 18 (246) | 1,000,543,756 | 24,051,369 | 83,706,672 | 67,280,866 | 7 | 33.85 |
| APSI_hap2_v1 | 67 (8,637) | 934,744,333 | 5,212 | 89,073,602 | 52,409,407 | 7 | 34.26 |
| APSI_hap2_v2 | 98 (301) | 1,050,666,308 | 12,623 | 96,196,788 | 61,710,375 | 7 | 33.86 |
| Chromosomes | 18 (221) | 1,043,460,636 | 27,407,230 | 96,196,788 | 61,710,375 | 7 | 33.86 |

1. Scaffold counts. Contig counts in brackets

**Figure 1.**
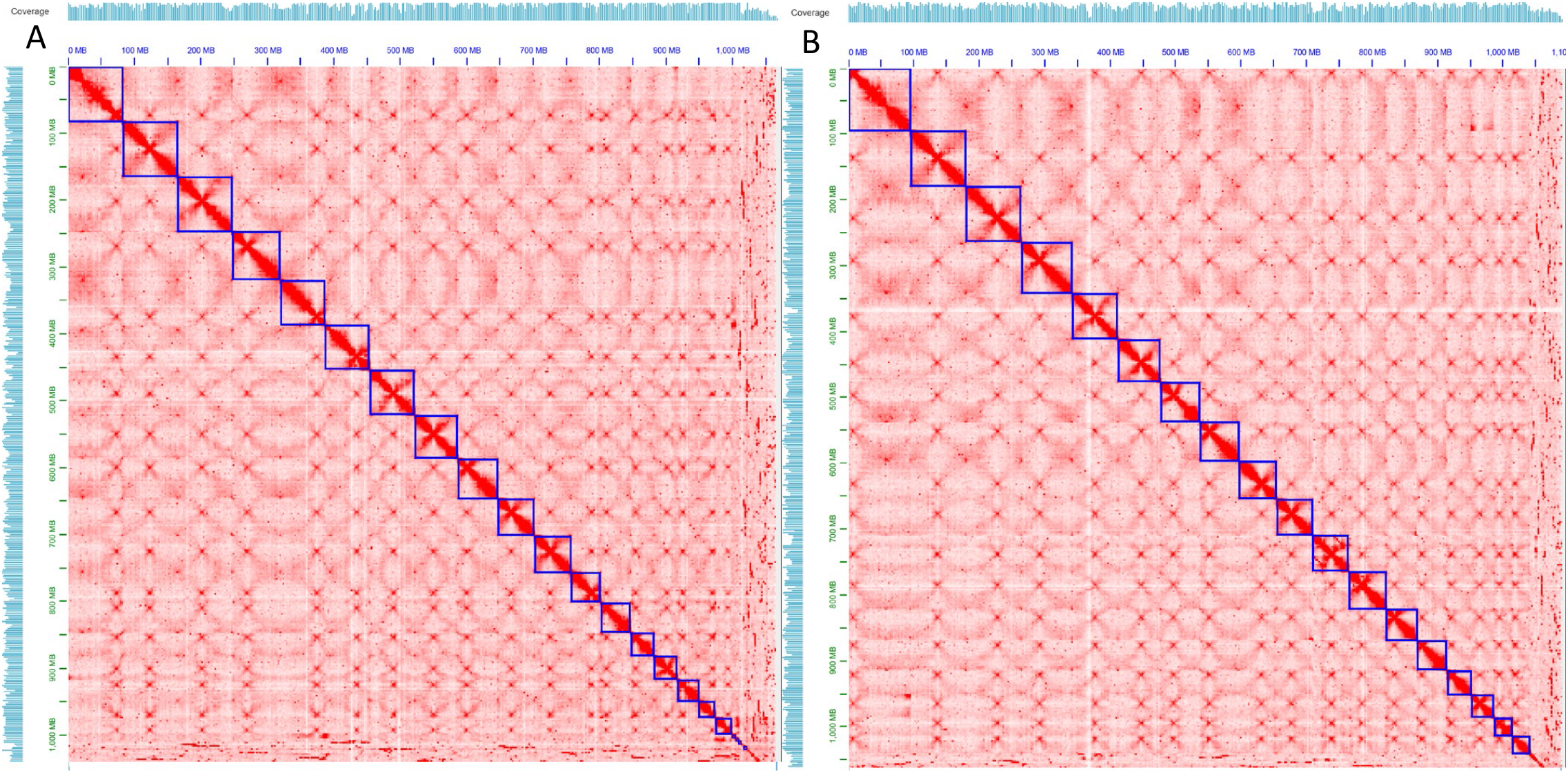
**Image from Juicebox visualisation software (v 1.11.08) of the scaffolded *Austropuccinia psidii* genome assembly. APSI_hap1 (A) and APSI_hap2 (B). Hi-C coverage indicted by pale blue bars, left and top. Blue squares indicate the predicted 18 chromosomes in order of size. Smaller contigs, including those assembled as mitochondrial DNA, are retained as non-scaffolded contigs.**

**Figure 2.**
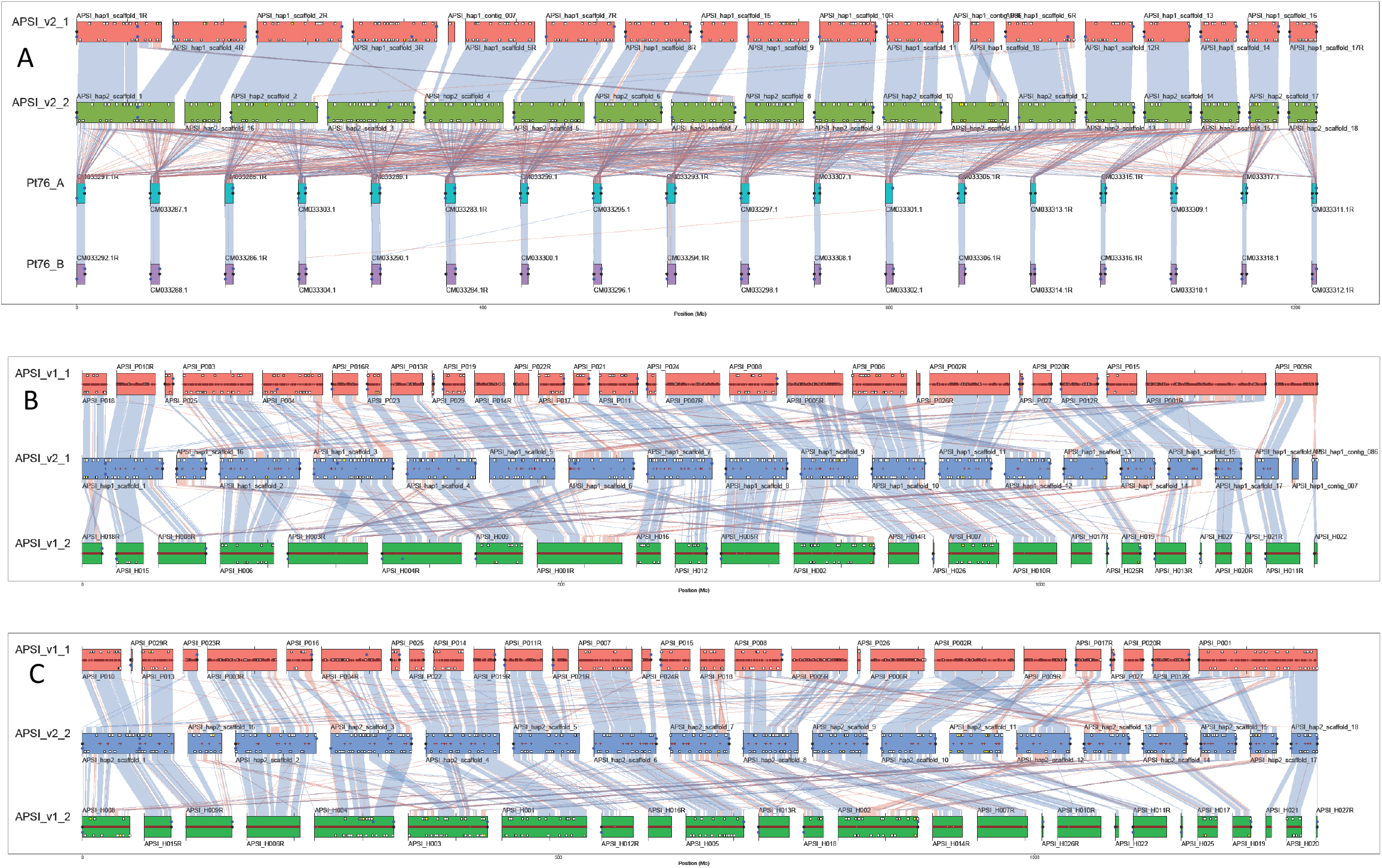
**ChromSyn BUSCO-derived synteny plots. Coloured blocks represent assembly scaffolds over 1Mbp for each genome. Synteny blocks of collinear “Complete” BUSCO genes link scaffolds from adjacent assemblies: blue, same strand; red, inverse strand. Filled circles mark telomere predictions from Diploidocus (black) and TIDK (blue). Assembly gaps are marked as dark red + signs. Predicted v1 *A. psidii* effector coding sequences (Tobias *et al*. 2021) are marked as white squares, or yellow when haplotype-specific. Genes on the forward strand are marked above the centre-line, with reverse strand genes below. (A) Pairs of phased haploid genomes for *A. psidii* v2 and wheat leaf rust chromosomes, Pt76_A and B (Duan *et al*. 2022). Assembly gaps not shown. (B) Improved APSI_v2_1 (hap1) assembly, flanked by both v1 haploid assemblies. (C) Improved APSI_v2_2 (hap2) assembly, flanked by both v1 haploid assemblies.**

### Improved nuclear haploid genome assemblies

A comparison of *A. psidii* genome versions one and two show that the new assemblies have longer minimum sequence lengths, better resolved N50 but smaller maximum length (Table 1). Although the scaffold numbers are higher, v2 is considerably more contiguous and has over 98% allocated to the 18 largest scaffolds of each haplotype (Table 1, Figure 2). Synteny and telomere analysis indicates that the long v1 scaffolds are likely to be mis-scaffolded contigs from multiple chromosomes (Figure 2). Synteny between the hap1 and hap2 assemblies of v2 is generally very high but indicates that a few mis-assemblies remain at the chromosome level, particularly where telomeres are not found at each end of the scaffold. The overall v2 BUSCO completeness of 92.0% is a slight improvement over 91.4% for v1. This is comparable to the overall BUSCO completeness scores for wheat leaf rust Pt76 of 91.0%, suggesting that the missing BUSCO genes might represent lineage-specific losses in rust fungi rather than missing data from the assemblies. The total genome size for v2 is very similar to v1, supporting previous observations about expansions of transposable element families. Curiously, DepthKopy analysis of scaffolds and 100 kb sliding windows (Figure 5) reveals a dominant read depth somewhere between *n* and *2n*. This is not seen to the same extent for mean read depth (data not shown) and may reflect that the transposable elements are not yet fully resolved, causing a minority of copies to attract an excess of mapped reads resulting in a genome-wide intergenic depletion of read depth.

### A complete and circularised *Austropuccinia psidii* mtDNA genome

In total, 9,964 HiFi reads (141.3 Mb) were mapped onto hap1 mtDNA regions and extracted for an initial draft mtDNA assembly. Subsequently, 7,192 HiFi reads (102.4 Mb) met the criteria of at least 10 kb and 99% of the read mapping to the draft mtDNA. These reads were assembled, polished and circularised into a contiguous 94,553 bp circular mtDNA genome (37.42% GC), APSIMTv2. The previously published mtDNA genome from the MF-1 isolate of *A. psidii* was assembled with a combination of sequence read-types (NC_044121.1) and is 93,299 bp (de Almeida *et al*. 2021). The end of NC_004121.1 consists of 154 bp of a TAAACA-rich repeat, which is 200 bp long in APSIMTv2. It is not clear whether this represents a misassembly in either mtDNA or variable mtDNA sequences in the *A. psidii* population. The extra 974 bp of APSIMTv2was searched against NCBInr using discontinuous megablast (Altschul *et al*. 1990) 21/03/22] but yielded no notable hits. However, a BLAST+ search against the nuclear genome revealed several exact matches and should be investigated further as a possible NUMT. The APSI_hap1_v2 genome (NCBI accession JALGQZ000000000) incorporates the fully circularised mitochondrial DNA sequence as the final scaffold.

### Effectors located on chromosomes

We mapped 617 and 616 of the v1 predicted effector coding sequences onto hap1 and hap2 respectively. Chromosome three has an apparent abundance of predicted effector genes, evident in Figure 4, numbering 68 and 67 respectively. Discrepancies in effector gene numbers, notably scaffold 1, 7, 8, 9 and 10, are likely to be issues with current mis-scaffolding, as can be determined by the synteny graphs in Figure 2. However, it is notable that 22 predicted effector genes present in hap1 are absent in hap2, and likewise 21 genes present in hap2 are absent in hap1. These findings suggest high levels of heterozygosity for these pathogenicity genes. We were able to determine a number of predicted effector genes within proximity of telomeric regions, known for fungal effector diversity (Gan *et al*. 2021), as presented in Figure DepthKopy analysis of the effector genes reveals a very similar predicted copy number distribution to “Complete” (single copy) BUSCO genes (Figure 3). An increase is observed in the proportion of effector genes with a predicted copy number of 0.5, which corresponds to the haploid *n* read depth rather than diploid *2n* depth, especially for the effector genes only found in hap2 (Figure 5). This could indicate hemizygous genes that are only found in a single haploid genome, although a false duplication in the assembly would give a similar signal. A similar but more pronounced signal from Duplicated BUSCO genes indicates that a small, but non-zero, number of false duplicates remain, with the level of BUSCO duplication also higher in hap2 (2.0%) than hap1 (1.2%). Many Duplicated BUSCO genes exhibit a normal *2n* depth profile, however, consistent with genuine lineage-specific duplications (Chen *et al*. 2022), particularly for hap1.

**Figure 3.**
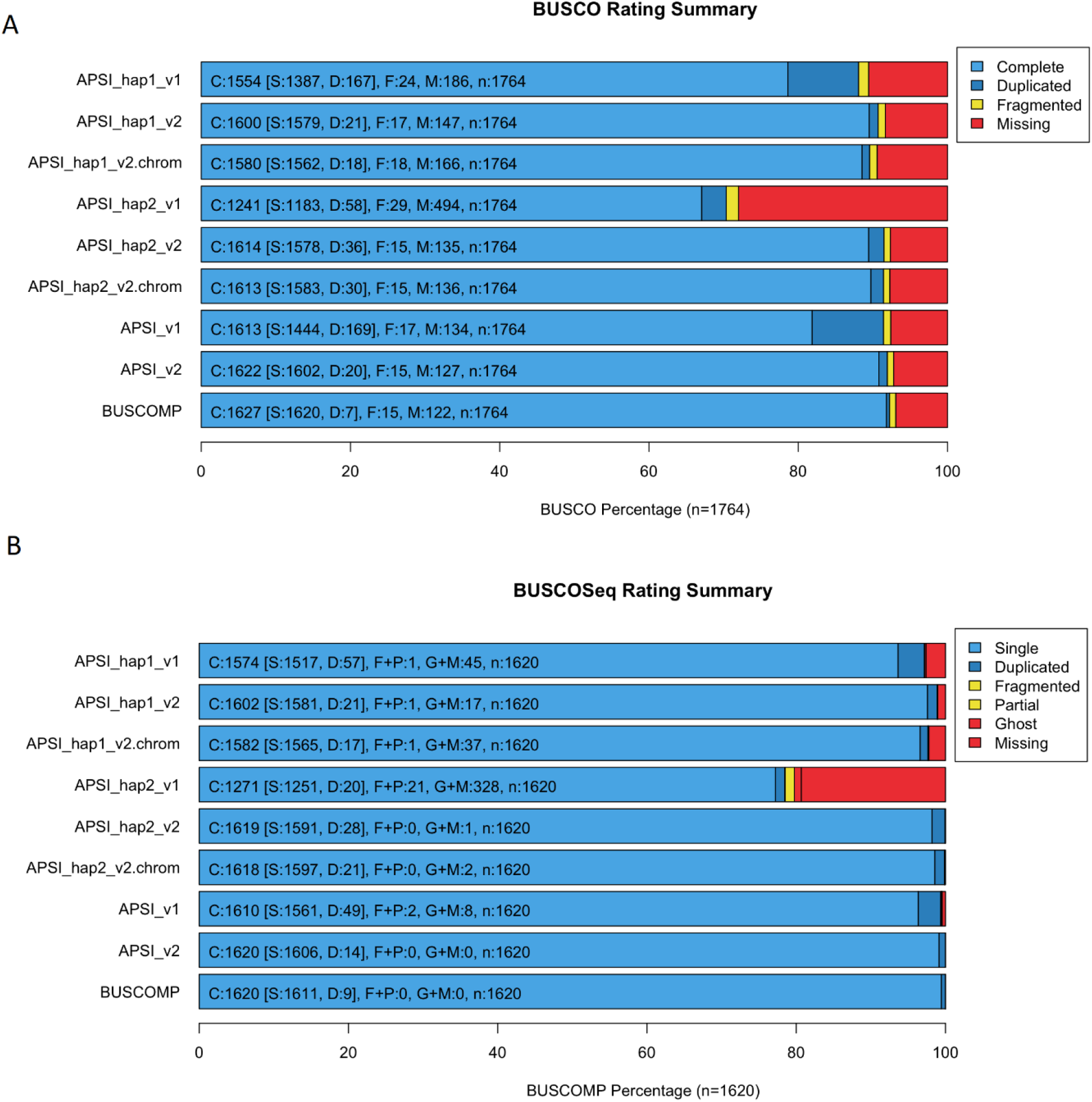
**BUSCOMP compilation of BUSCO v5.2.2 results for v1and v2of the *Austropuccinia psidii* assembly. chrom, chromosome scaffolds only; APSI_v1, compiled hap1_v1 and hap2_v1 results; APSI_v2, compiled hap1_v2.1 and hap2_v2.1 results; BUSCOMP, compiled results from all assemblies. (A) BUSCO ratings for 1764 basidiomycota_odb10 genes. (B) BUSCOMP ratings for 1620 genes rated as single-copy “Complete” in at least one genome.**

**Figure 4.**
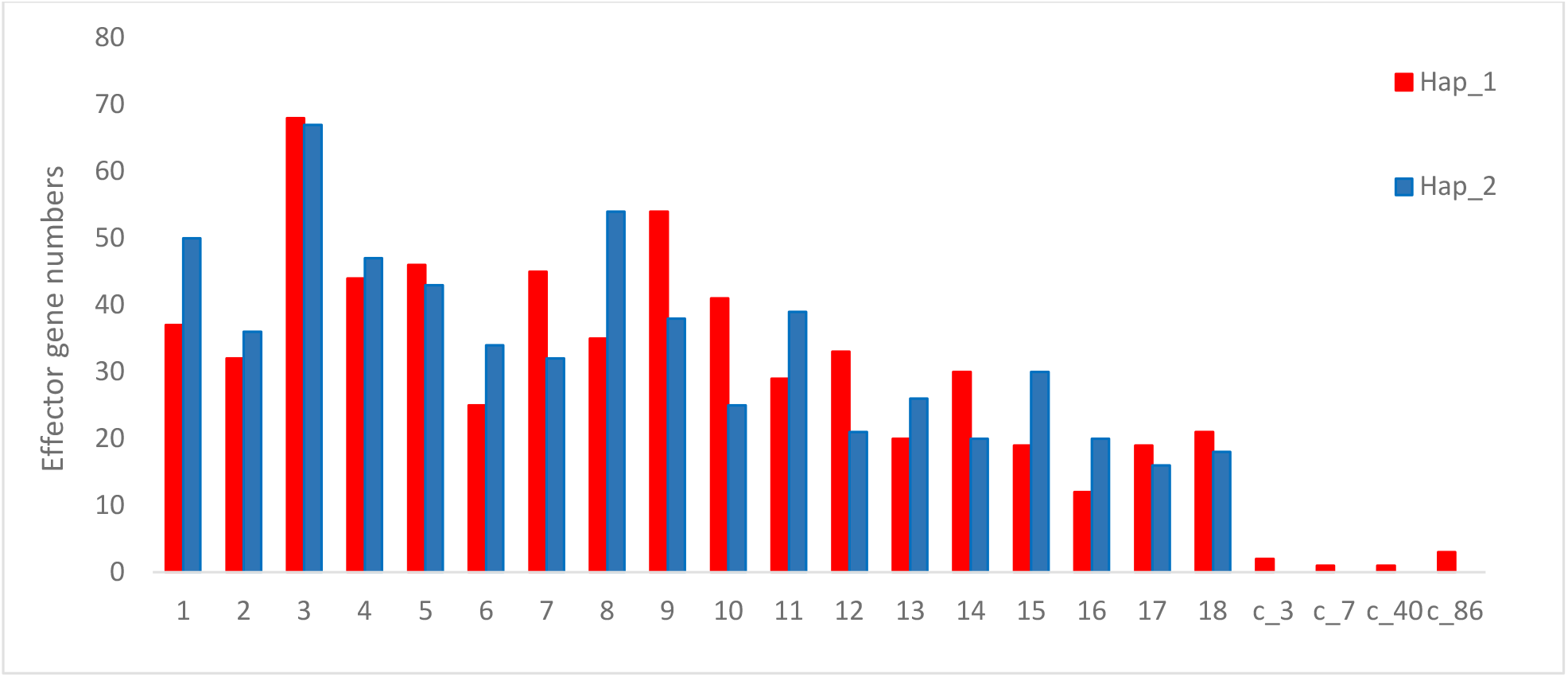
**Predicted effector genes, determined from annotated *A. psidii* v1 genome (Tobias *et al*. 2021), were mapped to chromosomes (x-axis) and contigs (denoted c_) of the two genome haplotypes (APSI_hap1 and APSI_hap2) with the largest numbers occurring on scaffold 3. Discrepancies in gene numbers between haplotypes, notably scaffold 1, 8 and 9, are likely to be issues with current mis scaffolding.**

**Figure 5.**
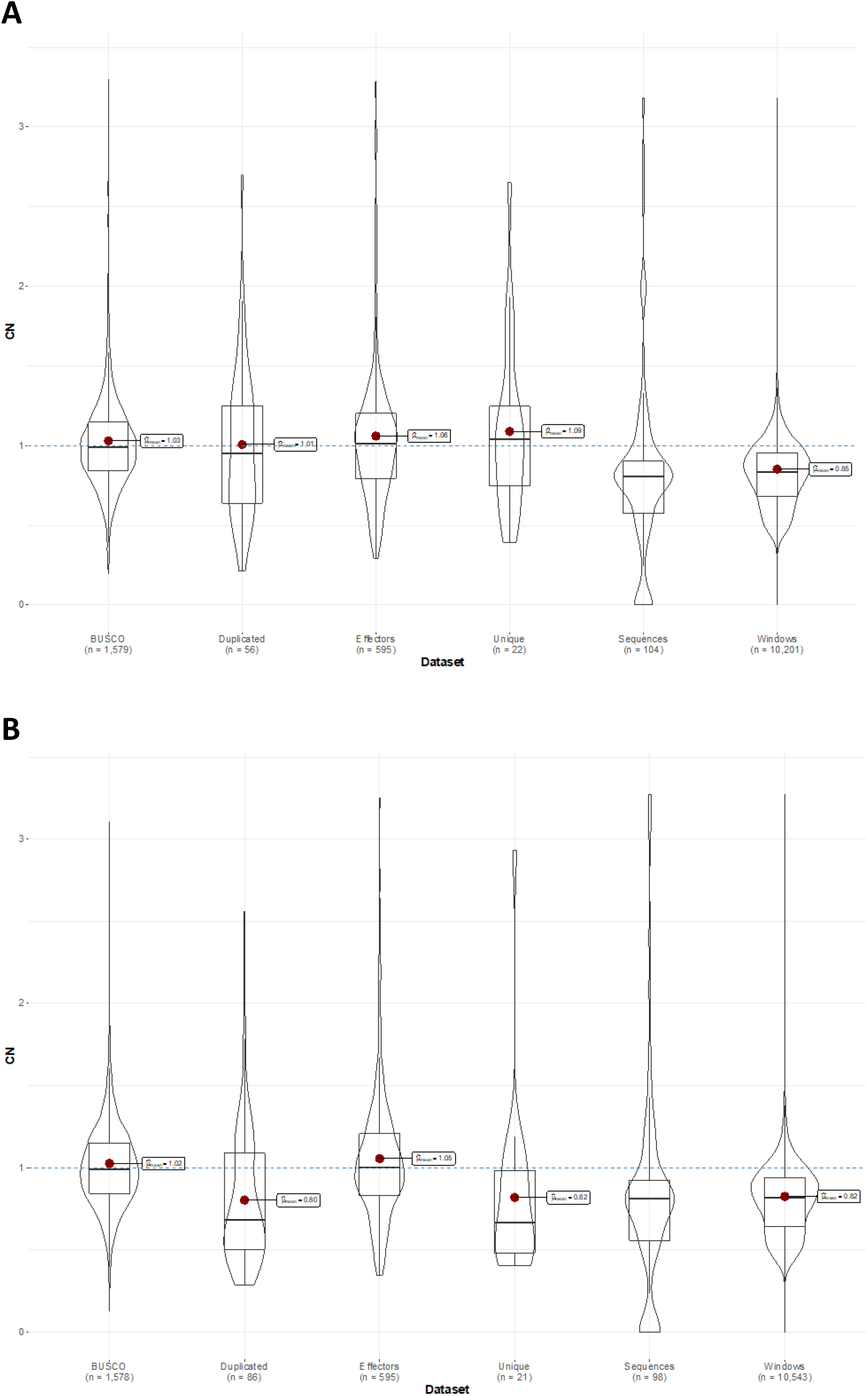
**DepthKopy predicted copy number plots for (A) hap1 and (B) hap2. BUSCO, “Complete” rated BUSCO genes; Duplicated, “Duplicated” rated BUSCO genes; Effectors, predicted effector protein genes found in both haploid genomes; Unique, predicted effector protein genes found in only one haploid genome; Sequences, assembly scaffold; Windows, 100 kb sliding wondows.**

## Discussion

### Rust chromosome numbers are highly conserved

Our research presents a semi-resolved genome ultra-structure for the pandemic biotype of *A. psidii*. The newly scaffolded genome validates the previous assembly findings regarding the size, extensive telomeric regions and intergenic expansions (Tobias *et al*. 2021). However we present a much improved assembly with high synteny of conserved genes between the haploid chromosomes, and compared to the genome for wheat leaf rust, *Puccinia triticina* (Pt76) (Duan *et al*. 2022) (Figure 2).

Here we show for the first time that *A. psidii* has 18 haploid chromosomes and that 12 of these have telomeres located at both ends, in both haploid genomes, indicating complete chromosomes. *Austropuccinia psidii* thereby conforms to the published karyotypes for other rust species (Boehm *et al*. 1992b; Boehm and Bushnell 1992). Early experiments to karyotype rust fungi under-represented chromosome numbers at between *n* = 3-6 (Valkoun and Bartoš 1974; Goddard 1976) due to the small physical size and difficulty of capturing the metaphase state in biotrophs. Later cytogenetic studies in several rust species including *Melampsora lini* (Boehm and Bushnell 1992) and *Puccinia graminis* f.sp. *tritici* (Boehm *et al*. 1992b) determined a consistent haploid chromosome number of *n*=18. These findings have been recently further confirmed with new technology that has enabled phasing and scaffolding of genomes for wheat leaf rust (Wu *et al*. 2021; Duan *et al*. 2022) and stripe rust (Schwessinger *et al*. 2018, 2022). While the genome for *A. psidii* conforms to these numbers, this finding was not assured due to the surprising 1 Gb size of the genome and its evolutionary lineage within the distantly related Sphaerophragmiaceae family (Beenken 2017). Despite the large diversity of rust-type fungi and evolutionary radiation prior to 85 million years ago (Aime *et al*. 2018), our results indicate that chromosome numbers are highly conserved. A key finding of Tobias et al. 2021 was that the genome size is largely driven by intergenic expansions of transposable elements (TE). In Figure 2, the *A. psidii* chromosome sizes relative to the much smaller wheat leaf rust (Pt76) chromosomes, are clearly visible.

### ChromSyn: a new tool for synteny analyses

Genome-wide synteny analysis of chromosome-level assemblies can be extremely useful for identifying possible scaffolding errors in phased haploid genomes (Figure 2A) or investigating chromosome evolution across species (Waters *et al*. 2021). Full genome alignments are computationally intensive, especially for species that are rich with repetitive sequences, such as *A. psidii*. Here, we demonstrate an efficient new tool for chromosome synteny analysis, Chromsyn, which utilises inferred synteny blocks derived from single-copy (“Complete”) BUSCO genes. BUSCO itself is a highly efficient tool for identifying orthologous regions in genomes and has the additional advantage of being routinely run for completeness assessment of genomes, eliminating the need for additional computation. Whilst local rearrangements between BUSCO genes be missed by ChromSyn, we demonstrate how the high-level synteny visualisation can be used to identify scaffolding issues within the previously published *A. psidii* genome. To further help identify scaffolding errors and chromosome completeness, ChromSyn can also plot telomere predictions from Diploidocus and/or tidk, as well as assembly gaps. Additional features, such as genes, can also be plotted, demonstrated here with predicted effector genes.

### A phased genome enables heterozygosity studies in *Austropuccinia psidii*

Predicted effector genes between the haploid genomes indicate high levels of heterozygosity, as previously determined for oat crown rust and stripe rust isolates (Miller *et al*. 2018; Schwessinger *et al*. 2020). Effectors are characterised as proteins that are secreted by plant pathogens to facilitate infection and manipulate hosts (Lo Presti *et al*. 2015; Sperschneider and Dodds 2022). They are important for pathogenicity on susceptible hosts, and for recognition by resistant hosts, and therefore drive strong evolutionarily selection in both pathogen and host. We compared the chromosome locations for predicted effector genes determined previously (Tobias *et al*. 2021) and can now show that variation occurs within haplotypes. We found evidence for 1,233 candidate effectors within the di-haploid genome, compared to ∼900 in oat crown rust (Miller *et al*. 2018) and 2,876 determined in wheat stripe rust (Schwessinger *et al*. 2022). Of these candidate effectors, we determined 595 shared gene homologs between haplotypes, but also that 22 and 21 gene models were exclusively present within hap1 and hap2 respectively. While numerical discrepancies of mapped genes by chromosome may be related to scaffolding issues, the overall mapped gene numbers per haplotype are likely to be accurate, based on the completeness of the genome assemblies. A complete annotation of the genome followed by downstream analyses will further clarify the haploid numbers and variation of effector genes in *A. psidii*.

A recent in-silico study into mating genes within *A. psidii* biotypes determined a tetrapolar mating type (Ferrarezi *et al*. 2022).The research determined the presence of a single copy of the pheromone receptors, STE3.2.2/4 and STE3.2.3 in all biotypes and two copies each of the linked homeodomain transcription factor genes, HD1 and HD2. On interrogation of the pandemic APSI_v2 phased genomes, it was determined that STE3.2.2/4 was exclusively present in hap2 and STE3.2.3 exclusively in hap1. This new finding determined the importance of these genes for mate compatibility and established, for the first time, that *A. psidii* use a tetrapolar mating strategy (Coelho *et al*. 2017). Previously, the presence of a single copy of a gene, within a well assembled but collapsed haploid genome, would indicate that these genes are likely to be highly conserved. Indeed, most genomes are currently published as collapsed haploid genomes. The data used for the mating gene analysis relied on the full phased genomes to determine an important biological truth that will facilitate a better understanding of pathogen population structures and sexual recombination potential.

### Telomere-capped full-length chromosomes confirm extensive telomeric size in *A. psidii*

Fifty-five of the putative chromosome ends had telomeres predicted, including thirty-six with Diploidocus length estimates (Table 2). Telomere length in basidiomycetes is species-specific (Lucía *et al*. 2010) but studies to date show a range between 100 and 300 bp (Pérez *et al*. 2009; Schwessinger *et al*. 2020; Sperschneider *et al*. 2020). We clarified that the telomeric repeats in *A. psidii* are much longer than other assembled fungal genomes, with over half (19/36) telomeres annotated with Diploidocus exceeding 5 kb. ChromSyn visualisation has also identified some anomalies present where telomeric repeats occur within chromosomes in a pattern that suggests some mis-scaffolding has occurred (Figure 2). Future work will further improve the chromosome-level scaffolding with an anticipated increase in the number of full-length chromosome scaffolds with telomeres at each end.

**Table 2.**
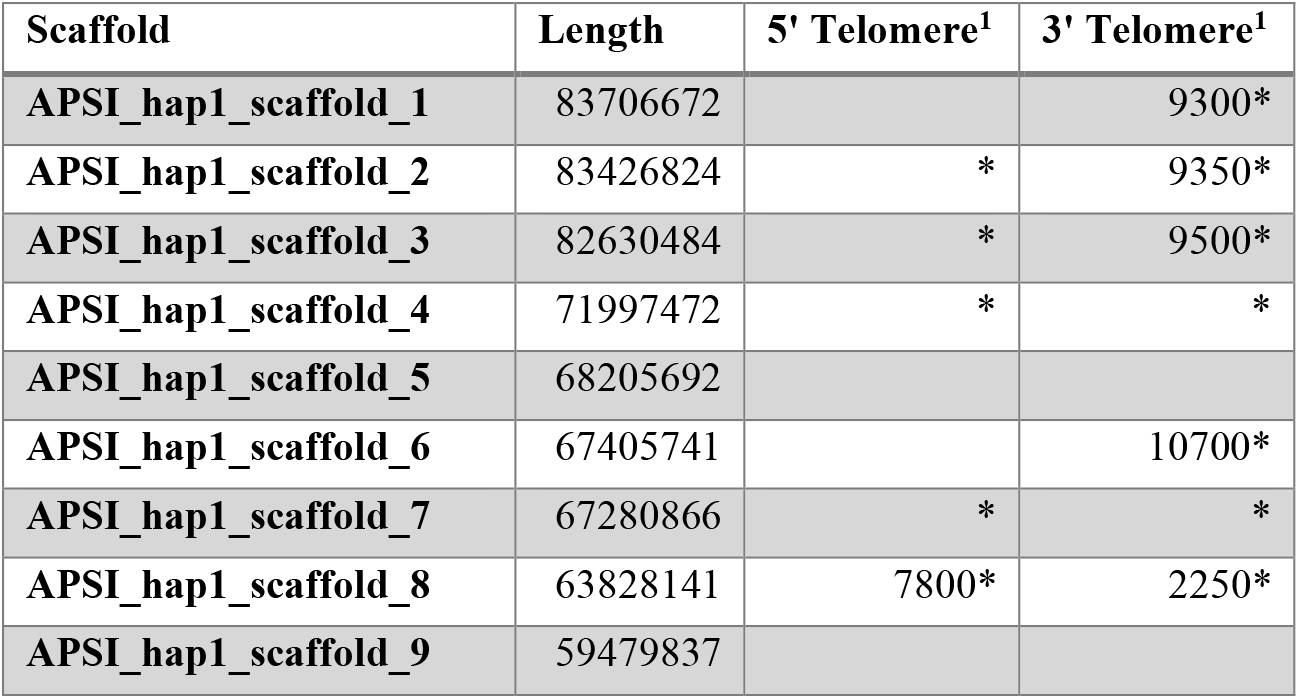
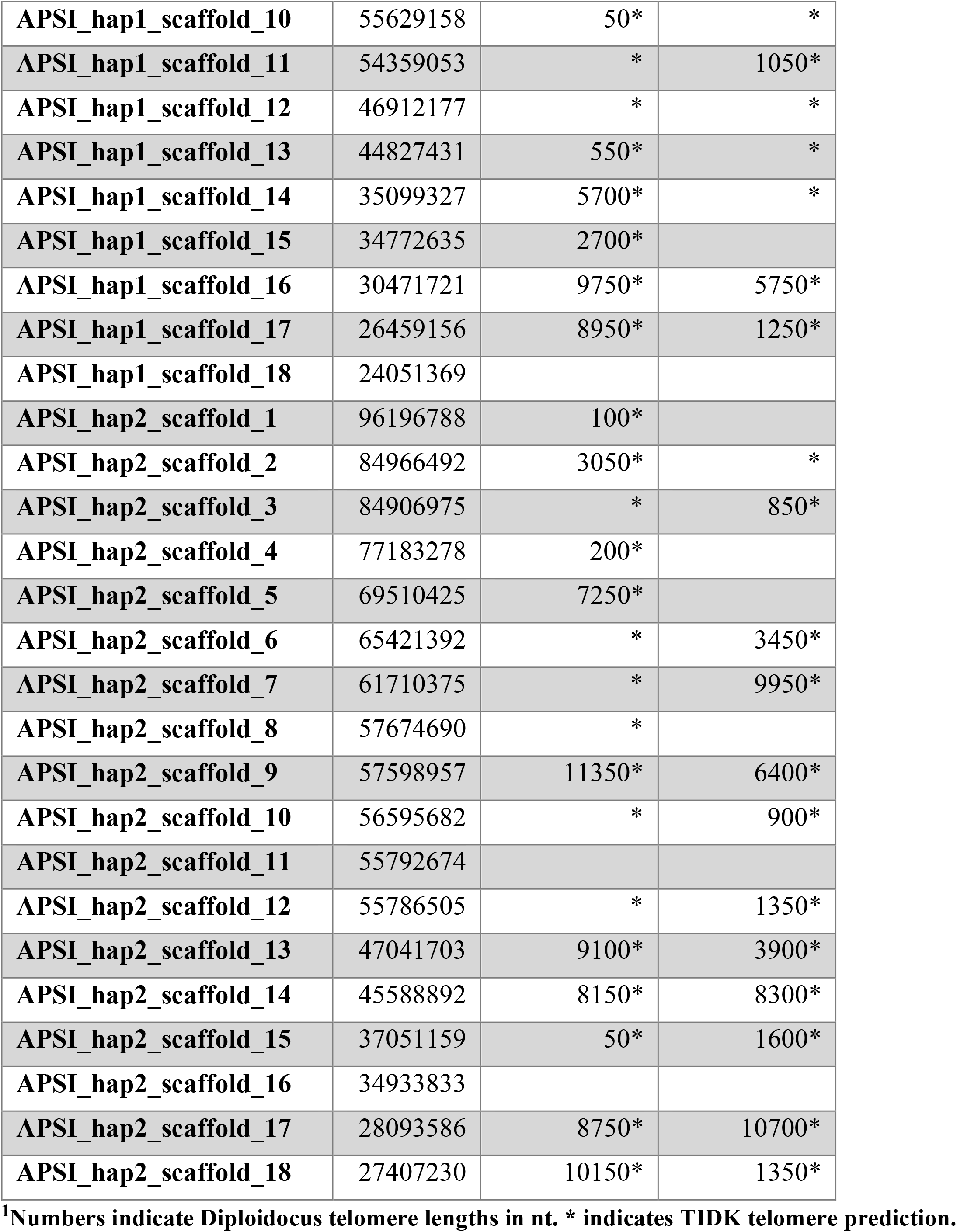
Diploidocus and TIDK telomere predictions for main chromosome scaffolds.

| Scaffold | Length | 5' Telomere<sup>1</sup> | 3' Telomere<sup>1</sup> |
| --- | --- | --- | --- |
| APSI_hap1_scaffold_1 | 83706672 |  | 9300* |
| APSI_hap1_scaffold_2 | 83426824 | * | 9350* |
| APSI_hap1_scaffold_3 | 82630484 | * | 9500* |
| APSI_hap1_scaffold_4 | 71997472 | * | * |
| APSI_hap1_scaffold_5 | 68205692 |  |  |
| APSI_hap1_scaffold_6 | 67405741 |  | 10700* |
| APSI_hap1_scaffold_7 | 67280866 | * | * |
| APSI_hap1_scaffold_8 | 63828141 | 7800* | 2250* |
| APSI_hap1_scaffold_9 | 59479837 |  |  |
| APSI_hap1_scaffold_10 | 55629158 | 50* | * |
| APSI_hap1_scaffold_11 | 54359053 | * | 1050* |
| APSI_hap1_scaffold_12 | 46912177 | * | * |
| APSI_hap1_scaffold_13 | 44827431 | 550* | * |
| APSI_hap1_scaffold_14 | 35099327 | 5700* | * |
| APSI_hap1_scaffold_15 | 34772635 | 2700* |  |
| APSI_hap1_scaffold_16 | 30471721 | 9750* | 5750* |
| APSI_hap1_scaffold_17 | 26459156 | 8950* | 1250* |
| APSI_hap1_scaffold_18 | 24051369 |  |  |
| APSI_hap2_scaffold_1 | 96196788 | 100* |  |
| APSI_hap2_scaffold_2 | 84966492 | 3050* | * |
| APSI_hap2_scaffold_3 | 84906975 | * | 850* |
| APSI_hap2_scaffold_4 | 77183278 | 200* |  |
| APSI_hap2_scaffold_5 | 69510425 | 7250* |  |
| APSI_hap2_scaffold_6 | 65421392 | * | 3450* |
| APSI_hap2_scaffold_7 | 61710375 | * | 9950* |
| APSI_hap2_scaffold_8 | 57674690 | * |  |
| APSI_hap2_scaffold_9 | 57598957 | 11350* | 6400* |
| APSI_hap2_scaffold_10 | 56595682 | * | 900* |
| APSI_hap2_scaffold_11 | 55792674 |  |  |
| APSI_hap2_scaffold_12 | 55786505 | * | 1350* |
| APSI_hap2_scaffold_13 | 47041703 | 9100* | 3900* |
| APSI_hap2_scaffold_14 | 45588892 | 8150* | 8300* |
| APSI_hap2_scaffold_15 | 37051159 | 50* | 1600* |
| APSI_hap2_scaffold_16 | 34933833 |  |  |
| APSI_hap2_scaffold_17 | 28093586 | 8750* | 10700* |
| APSI_hap2_scaffold_18 | 27407230 | 10150* | 1350* |

<sup>1</sup>Numbers indicate *Diploidocus* telomere lengths in nt. \* indicates TIDK telomere prediction.

## Conclusion

We have assembled a phased chromosome-level genome for the pandemic biotype of *Austropuccinia psidii*. Our improved genome has clarified the diploid chromosome numbers at 36, and we show that 24 of these are full-length with both 5’ and 3’ telomeric sequences. Many serious knowledge gaps around pathogenicity cannot be determined without investigations into both parental alleles. It is hoped that this resource will enable progress into many of the unanswered questions around host susceptibility, fungal mating strategies, effector gene complements and heterozygosity within this dikaryotic tree pathogen.

## Acknowledgements

Grant funds supporting this research came from the Australian Flora Foundation. The authors acknowledge the Sydney Informatics Hub and the University of Sydney’s High-Performance Computing cluster for providing the computing resources that have contributed to the research results reported within this paper.

## Author contributions

PAT initiated and led the research, sourced grant funding, assembled and scaffolded the genome, organised data submission, wrote much of manuscript. RJE assembled the mitochondrial genome, developed software that was used to curate the genome data and run analyses, prepared figures, contributed to the methods and results of the manuscript writing. CD extracted HMW DNA. R.F.P. supported the study, contributed to the manuscript, and provided the spore samples.

## Data availability

This Whole Genome Shotgun project for *Austropuccinia psidii* Au3-C622-A115012 has been deposited at DDBJ/ENA/GenBank under the accessions JALGQZ000000000 and JALGRA000000000. Genomes are accessible here: 10.5281/zenodo.6476632. The complete and circularised mitochondrial sequence can be found as a final scaffold in the APSI_hap1_v2 genome: JALGQZ000000000. Raw data is available at the PRJNA810573/ PRJNA81057

## Notes

https://zenodo.org/record/6476632#.YmIuWNpBxPY

### Competing Interest Statement

The authors have declared no competing interest.

